# Dilution and titration of cell-cycle regulators may control cell size in budding yeast

**DOI:** 10.1101/300160

**Authors:** Frank S. Heldt, Reece Lunstone, John J. Tyson, Béla Novák

## Abstract

The size of a cell sets the scale for all biochemical processes within it, thereby affecting cellular fitness and survival. Hence, cell size needs to be kept within certain limits and relatively constant over multiple generations. However, how cells measure their size and use this information to regulate growth and division remains controversial. Here, we present two mechanistic mathematical models of the budding yeast (*S. cerevisiae*) cell cycle to investigate competing hypotheses on size control: inhibitor dilution and titration of nuclear sites. Our results suggest that an inhibitor-dilution mechanism, in which cell growth dilutes the transcriptional inhibitor Whi5 against the constant activator Cln3, can facilitate size homeostasis. This is achieved by utilising a positive feedback loop to establish a fixed size threshold for the START transition, which efficiently couples cell growth to cell cycle progression. Yet, we show that inhibitor dilution cannot reproduce the size of mutants that alter the cell’s overall ploidy and *WHI5* gene copy number. By contrast, size control through titration of Cln3 against a constant number of genomic binding sites for the transcription factor SBF recapitulates both size homeostasis and the size of these mutant strains. Moreover, this model produces an imperfect ‘sizer’ behaviour in G1 and a ‘timer’ in S/G2/M, which combine to yield an ‘adder’ over the whole cell cycle; an observation recently made in experiments. Hence, our model connects these phenomenological data with the molecular details of the cell cycle, providing a systems-level perspective of budding yeast size control.

## Introduction

Balanced growth of proliferating cells requires some coordination between the increasing size of a growing cell and its probability of undergoing DNA synthesis and division. In particular, the average time between two successive cell divisions must allow for a doubling in cell mass (or volume, which we will use interchangeably in the following). Any systematic deviation from this balance would lead to progressive changes in size over consecutive generations, eventually leading to the breakdown of biochemical processes. However, despite mounting evidence for active size control in various cell types and across different organisms [1], if and how cells measure their size and relay this information to the cell cycle remains controversial [2].

An elegant way to coordinate cell division and growth is to restrict passage through a certain cell cycle stage to cells larger than a particular target size [1]. Such ‘size checkpoints’ have been proposed to underlie size control at the START transition in budding yeast [3–5], and at the G2/M transition in fission yeast [6–8] and slime mould plasmodia [9–11]. The critical size required to pass these transitions depends, among other things, on the ploidy of the cell and its nutritional status [2]. To establish a size checkpoint, cells need to generate a size-dependent biochemical signal. Yet, most cellular macromolecules increase in abundance proportionally to cell volume, so that their concentration remains constant and the reactions they are involved in are independent of size [12]. Several proteins that defy this general rule have been indicated in size control. The mitotic activator Cdc25, for instance, increases in concentration with size in fission yeast [8], while Whi5, an inhibitor of START in budding yeast, is diluted by cell growth [13]. This suggest a general mechanism, in which size control emerges from the interplay between size-dependent and size-independent cell cycle regulators. Here, we study this intriguing possibility, focusing on the budding yeast cell cycle.

The budding yeast *Saccharomyces cerevisiae* divides asymmetrically, with size control mainly operating in the new-born daughter cell when it commits to enter the cell cycle anew at the START transition [3–5]. Passage through START is driven by activation of the transcription factor SBF [14]. In early G1-phase, before START, SBF is kept inactive by its stoichiometric inhibitor Whi5 [15,16]. To enter the cell cycle, the cyclin-dependent kinase Cdk1 (encoded by the *CDC28* gene) in conjunction with its regulatory binding partner Cln3 phosphorylates Whi5, which partially liberates SBF from inhibition and induces the synthesis of other G1 cyclins (Cln1 and Cln2). Cln 1/2:Cdk1 complexes then accelerate the phosphorylation of Whi5 and activation of SBF, thereby promoting the START transition [15–17]. Recent experiments show that during G1 the concentration of Cln3, the activator of START, is constant, while the concentration of Whi5 decreases, suggesting that an inhibitor-dilution mechanism facilitates size control [13]. However, previous theoretical considerations and experimental data suggested a different mechanism based on the titration of an activator that increases in molecule number during growth – as would be the case if its concentration is kept constant – against a fixed number of nuclear sites [18–20].

To test these hypotheses, we developed a mechanistic mathematical model of the budding yeast cell cycle. At its core, the model comprises a simple description of gene expression in which both size-dependent and size-independent synthesis of proteins emerge seamlessly for a differential affinity of genes for ‘transcription machinery’. This allows size-dependent proteins to maintain a fixed concentration during growth without the need for complex, gene-specific regulation and for size-independent proteins to maintain a fixed number of molecules per cell. Together, such size-dependent and -independent proteins can generate size-dependent biochemical signals for progression through the cell cycle. Using this model, we show that an inhibitor-dilution mechanism can facilitate size homeostasis and correctly account for changes in protein synthesis observed in experiments that perturb the number of gene copies of cell cycle regulators as well as the overall ploidy of the cell. However, the model fails to reproduce changes in cell size seen in some of these mutants. Intriguingly, a combination of inhibitor dilution and the titration of an activator against genomic sites correctly recapitulates these changes in cell size. Such a model also produces cell size patterns consistent with a ‘sizer’ mechanism in G1 and a ‘timer’ period comprising S, G2 and M-phase, which combine to yield an adder-type behaviour over the entire cell cycle; an observation recently made experimentally [21]. Hence, our model unites various experimental findings that were previously thought incompatible.

## Results

### A model for size-dependent and -independent protein expression

Experimental evidence suggests that size control emerges from the interplay of regulatory proteins whose synthesis rates depend on cell size and their size-independent counterparts [8,13]. To simulate the expression of such proteins we propose a simple mathematical model based on the differential binding of transcription machinery (TM) to genes (Fig. 1A). We model cell growth by assuming that components of the TM are themselves synthesised from size-dependent genes, which makes the production of TM autocatalytic, and, furthermore, that products of size-dependent genes control the increase in cell volume. These simple assumptions result in an exponential rise in both the amount of TM and cell size over time (Fig. 1B), as is characteristic for budding yeast both in single cells and at the population level [2,21].

**Figure 1.**
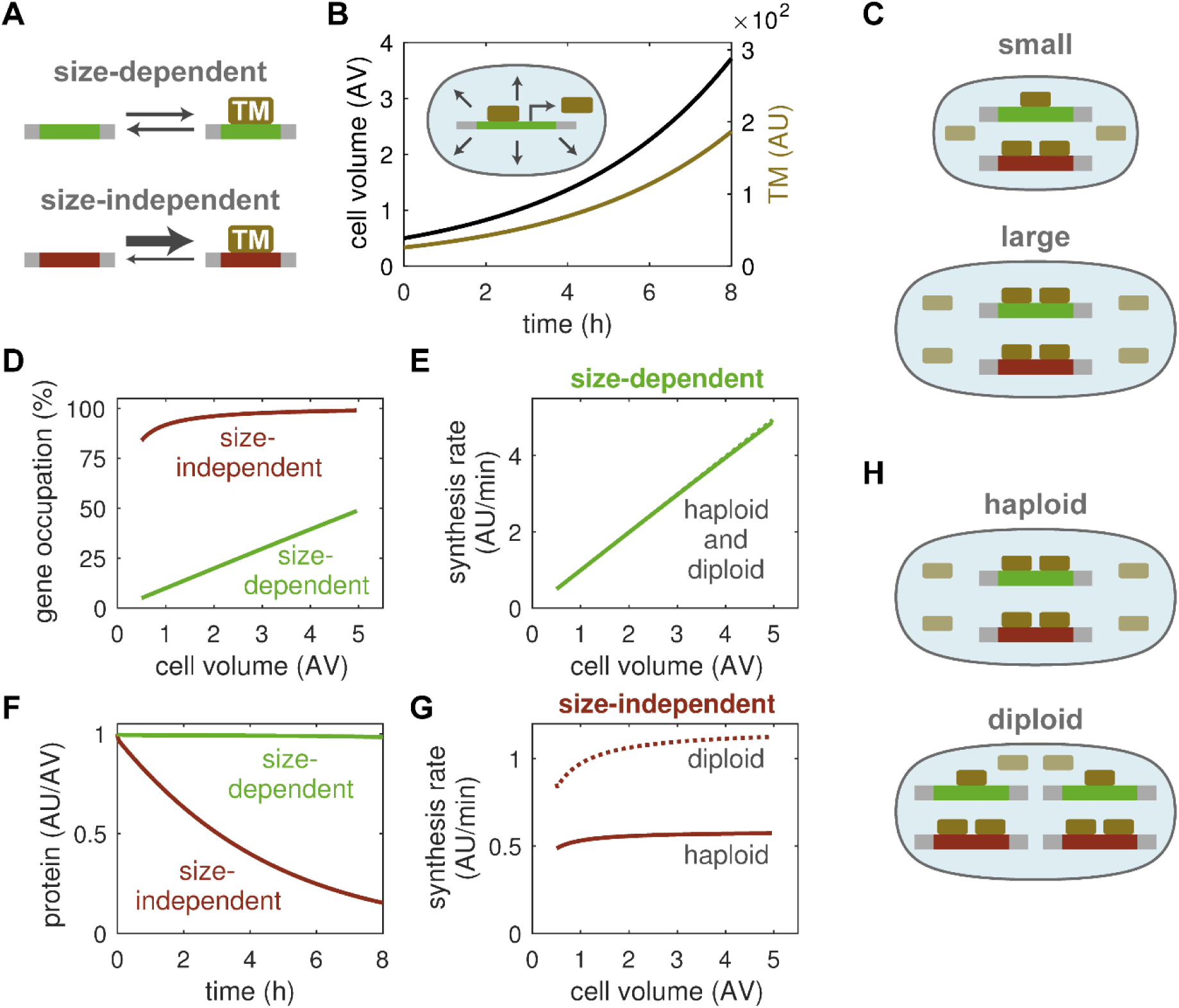
A model for size-dependent and -independent gene expression. (**A**) Transcription machinery (TM) binds with high affinity to the genes of size-independent proteins (bottom), while binding to the genes of size-dependent proteins is weaker (top). (**B**) Cell size and amount of TM in a model where growth and TM synthesis are controlled by size-dependent genes. (**C**) The amount of TM increases with cell size, which benefits the expression of size-dependent genes, while size-independent genes are saturated early, at low levels of TM. (**D**) Gene occupation by TM over cell volume. (**E, G**) Protein synthesis rates from size-dependent (E) and size-independent (G) genes in haploid (solid) and diploid cells (dashed). Curves in E overlap. (**F**) Concentration of a size-dependent and a size-independent protein over time in a growing cell. (**H**) Copy number affects the expression of size-independent genes as they efficiently compete for TM. Size-dependent genes split the TM among themselves. AU, arbitrary unit of number of molecules; AV, arbitrary unit of cell volume; AU/AV, arbitrary unit of concentration.

We note that the accumulation of TM in our simulations is compatible with experimental data on RNA polymerase II, which has been implicated in global transcriptional control [22]. Moreover, cell growth in the model depends on proteins that are themselves made by TM, which naturally leads to a direct proportionality between cell volume and transcriptional capacity. More precisely, as cells produce more and more TM their volume growth rate increases by the same extent, such that the number of TM molecules per unit cell volume remains constant. The fact that larger cells contain more TM translates into an increased occupation of size-dependent genes by TM, while size-independent genes are already fully occupied in small cells due to their high affinity for TM (Fig. 1C,D). Consequently, the transcriptional output from size-dependent genes increases with cell size (Fig. 1E), allowing their proteins to maintain a constant concentration during exponential cell growth (Fig. 1F). By contrast, expression from size-independent genes remains almost constant (Fig. 1G), such that their proteins are diluted by cell growth (Fig. 1F). Note that this basic size-related regulation could superimpose on other forms of gene expression control (e.g. cell cycle-dependent expression patterns).

Our gene expression model also predicts that an increase in gene copy number strongly affects the expression of size-independent proteins (Fig. 1G), but not of their size-dependent counterparts (Fig. 1E). This is because high-affinity, size-independent genes compete more efficiently for the shared pool of TM, such that an increase in their copy number directly translates into an increase in protein synthesis. Size-dependent genes instead split the available TM among themselves (Fig. 1H). In summary, our model uses a simple mechanism to explain why size-independent proteins are diluted by cell growth, whereas size-dependent proteins keep a constant concentration, without the need for complex, gene-specific regulation.

### Dilution of Whi5 can establish size control

Next, we asked whether the differential expression of cell cycle regulators according to the above model would allow budding yeast cells to control their size. In budding yeast, size control acts at START [3–5], where cells commit to cell cycle entry. Hence, we developed a cell cycle model centred on this transition (Fig. 2A). In this model, passage through START is facilitated by the activation of SBF, which is opposed by the stoichiometric inhibitor Whi5. Through the phosphorylation of Whi5, Cln3 liberates SBF from inhibition, thus driving cell cycle entry (Fig. S1A). Based on experimental observations [13], we assume that Whi5 is a size-independent gene, while all other proteins in our model are size-dependent. Consequently, cell growth in G1 dilutes the inhibitor of START, Whi5, while the activator Cln3 is maintained at constant concentration (Fig. 2B), as has been observed experimentally [13]. Our model shows that this inhibitor-dilution mechanism can establish a size threshold for START, where SBF is relieved from Whi5 inhibition only after sufficient growth has occurred (Fig. 2C). This transition is rapid and switch-like because of positive feedback via Cln1 and Cln2, which are expressed in response to SBF activation and further phosphorylate Whi5 [23,24]. The positive feedback loop creates a bistable switch, which implements the threshold response to graded changes in Whi5 concentration caused by cell volume growth, providing a sensitive size-sensing mechanism. After START has been passed, growth is restricted to the bud [4], and it continues until the end of the cycle, when the degradation of Clb1 and Clb2 initiates the separation of mother and daughter cell (Fig. 2D). Intriguingly, our model readily shows size homeostasis over multiple generations (Fig. 2D, lower panel). In particular, daughter cells, which we follow in our simulations because they show strong size control, reach the same size as their mothers, suggesting that Whi5 dilution can indeed couple cell division to cell growth.

**Figure 2.**
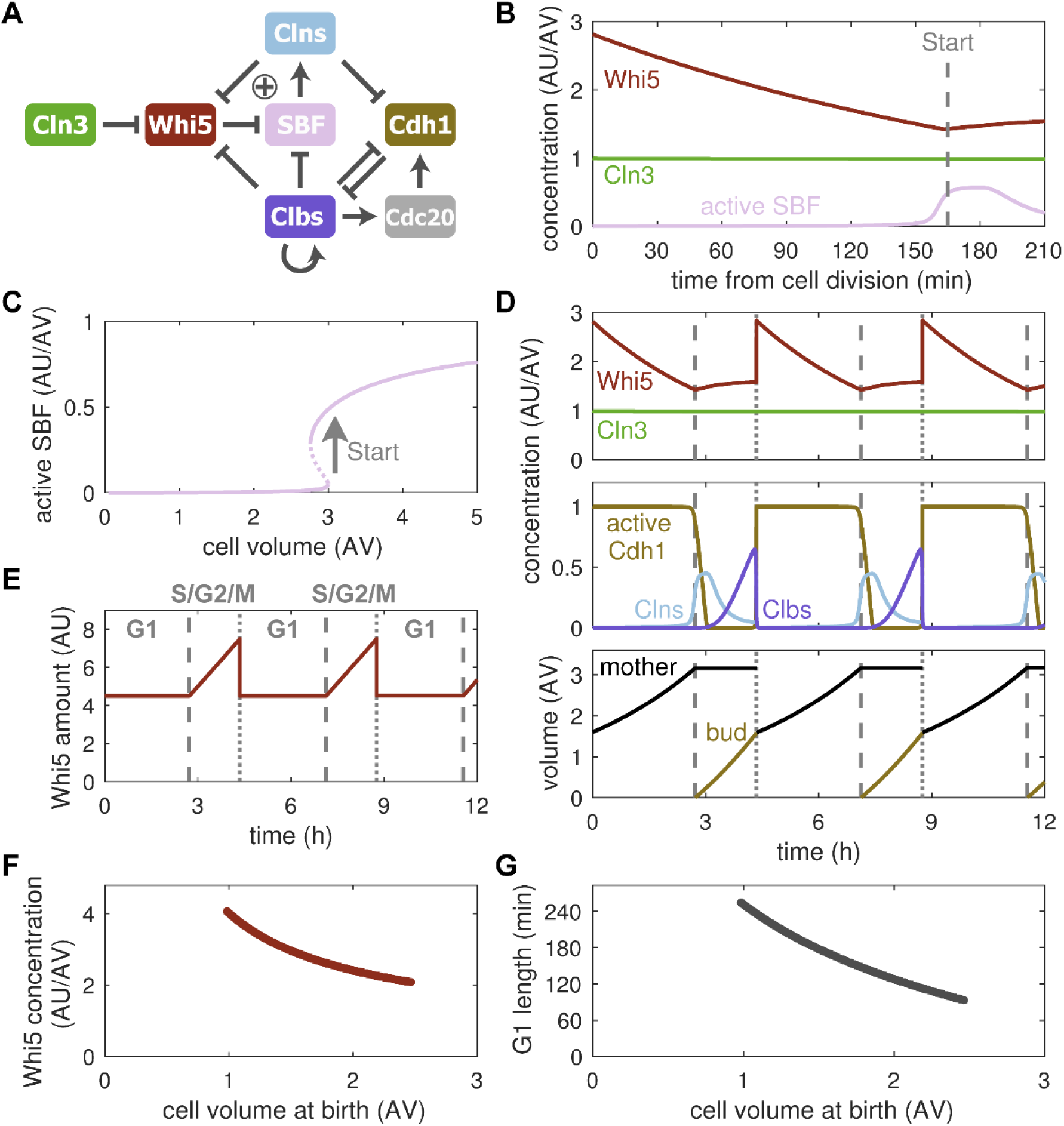
Inhibitor dilution allows for size homeostasis. (**A**) Influence diagram of the cell cycle model. Barbed edges indicate ‘activation’, blunt edges indicate ‘inhibition’, plus sign indicates positive feedback at START. The synthesis of all proteins except Whi5 is assumed to be size-dependent. (**B**) Concentration of the size-independent inhibitor of START, Whi5, and the size-dependent activator, Cln3, in G1-phase. Activation of SBF marks the onset of START (dashed line). (**C**) Stable (solid) and unstable (dashed) steady states of active SBF with respect to cell volume. Arrow indicates START transition. (**D**) Concentrations of cell cycle regulators (top and middle) and cell volume (bottom) over multiple generations. Dashed and dotted vertical lines mark START and cell division, respectively. Model follows the daughter cell (bud) after each division. (**E**) Simulated amount of Whi5 over multiple cell cycles. The G1-phase and S/G2/M period are indicated. (**F**) Correlation between Whi5 concentration and cell size at birth. Cell size was varied by changing the specific growth rate (see Supplementary Text for details). (**G**) Correlation between G1 length and cell size at birth.

In order to actively regulate cell size, i.e., to reduce size differences between cells, the inhibitor-dilution model requires that larger than average cells are born with lower than average Whi5 concentration so that they progress faster through G1, while smaller than average cells have higher Whi5 concentration, which gives them more time to grow. It has been proposed that this negative correlation between cell size at birth and Whi5 concentration results from the synthesis of a fixed amount of Whi5 during a period of fixed duration, which encompasses S-, G2- and M-phases in budding yeast [13]. By design our model accounts for this synthesis pattern, restricting Whi5 synthesis to the post-START period (Fig. 2E). We find that new-born cells do indeed exhibit a size-dependent Whi5 concentration (Fig. 2F). This allows for the adjustment of G1 duration to a cell’s birth size (Fig. 2G). In summary, our model demonstrates that size-independent synthesis of Whi5 and its dilution during G1 can allow cells to maintain their size over multiple generations by creating a cell-size threshold for START. Furthermore, the synthesis of a fixed amount of Whi5 per cell cycle can adjust for size differences by tuning G1 duration.

### Inhibitor-dilution model fails to reproduce all ploidy effects

To further explore the model’s ability to reproduce size control, we compared it to experiments that vary the copy number of *CLN3* and *WHI5*, as well as the cell’s overall ploidy [13]. These data were originally used to prove that Whi5’s synthesis rate is independent of cell size, while Cln3’s synthesis rate increases in larger cells ([13] and Fig. 3A). These experiments also highlight that Whi5 synthesis is largely independent of ploidy, with only a slight decrease seen between haploid and diploid cells that harbour the same number of *WHI5*copies (Fig. 3A, left panel). Yet, when the copy number of its gene is doubled, the Whi5 synthesis rate changes proportionally. Cln3 expression, in contrast, does change with ploidy, i.e., the slope of the synthesis rate decreases in diploid cells with one copy of *CLN3* compared to their haploid counterparts (Fig. 3A, right panel). However, an increase in *CLN3* copy number does not affect the Cln3 synthesis rate as long as the ratio between copy number and ploidy is kept constant. Crucially, diploid cells (with two copies each of *WHI5* and *CLN3*) were shown to be roughly twice the size of haploid cells (with one copy each of *WHI5* and *CLN3*).

**Figure 3.**
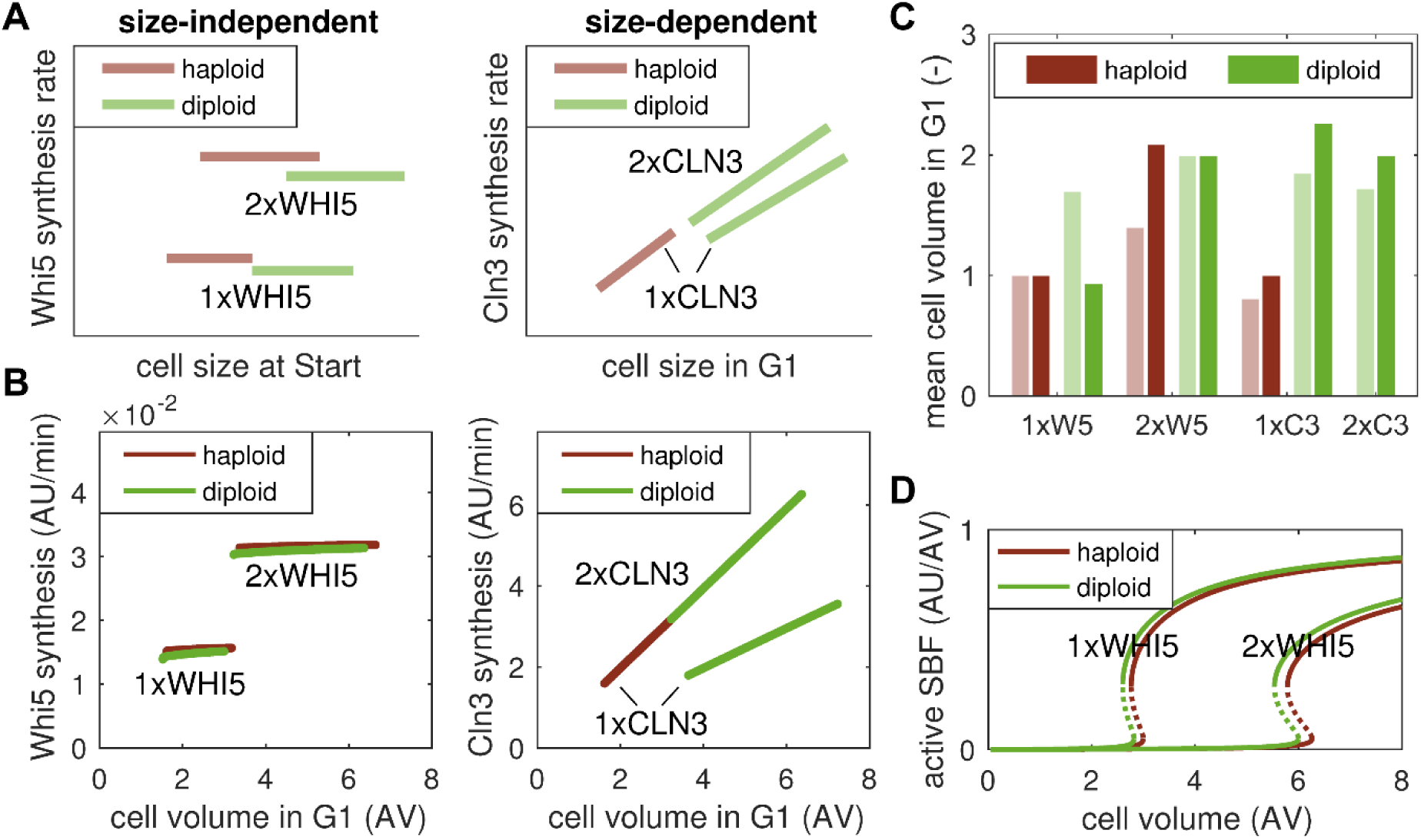
Inhibitor-dilution model fails to capture all ploidy effects on size. (**A**) Qualitative reproduction of experimental data in Fig. 2F and G of Ref. [13]. Graphs show the synthesis rates of Whi5 and Cln3 in haploid and diploid cells with the indicated copy number of *WHI5* and *CLN3* genes. (**B**) Simulation of Whi5 and Cln3 synthesis rates in haploid and diploid cells with the indicated copy number of *WHI5* and *CIN3.* (**C**) Mean cell volume in G1 for data in A (light bars) and simulations in B (dark bars). Values were normalized to haploid cells with one *WHI5* copy for each case. (**D**) Stable (solid) and unstable (dashed) steady states of active SBF with respect to cell volume for haploid and diploid cells with one and two *WHI5* copies.

We simulated these copy number mutants using the inhibitor-dilution model, which includes the features of gene expression shown in Fig. 1. The resulting simulations correctly predict the changes in protein synthesis rates for both Whi5 and Cln3 (Fig. 3B). In particular, they recapitulate the copy-number dependence of Whi5 synthesis rate and the copy-number-to-ploidy dependence of Cln3 synthesis rate. The model also correctly predicts the two-fold size increase between haploid and diploid cells. However, our simulations fail to reproduce the size increase observed between haploid and diploid cells with the same number of *WHI5* copies (Fig. 3C). More precisely, since protein synthesis rates for both Whi5 and Cln3 are similar in haploid and diploid cells with one *WHI5* copy (Fig. S2A), the model predicts a similar size threshold for START (Fig. 3D). In fact, considering the slight decrease in Whi5 synthesis rate observed in experiments [13], diploid cells with one *WHI5* should show a slight decrease in size compared to haploid cells according to our model. Reference [13] attributes the observed increase in cell size between haploid and diploid cells (with one or two copies of *WHI5*) to a delay in S/G2/M progression for diploid cells. Testing this hypothesis, we find that it only partially accounts for the observed size changes (Fig. S2B). In particular, diploid cells with one *WHI5* are predicted to be smaller than haploid cells with two *WHI5*, suggesting a larger influence of Whi5 synthesis rate than ploidy (Fig. S2C). However, in experiments the opposite is observed ([13] and Fig. 3A). Moreover, a delay in S/G2/M progression together with the observed increase in Whi5 synthesis rate would lead to a more than two-fold difference between haploid and diploid cells (Fig. S2C), in contradiction to experimental data [13]. Taken together, the inhibitor-dilution model thus correctly captures protein synthesis rates in copy number and ploidy mutants but fails to reproduce the observed size increase for some diploid cells.

### Titration of nuclear sites can account for ploidy effect

Previous theoretical and experimental studies attributed the effects of ploidy on cell size to an alternative control mechanism relying on the titration of a protein with constant concentration against a fixed number of nuclear sites [9–11,18–20]. In particular, it has been suggested that Cln3 is titrated against SBF bindings sites on the genome [20]. Based on these suggestions, we augmented the inhibitor-dilution model with a titration mechanism to test whether these two concepts can be brought into unison (Fig. 4A). In the pure inhibitor-dilution model, SBF, Whi5 and Cln3 interact in a strictly concentration-based manner (Fig. S1A). By contrast, the titration model assumes that SBF occupies a fixed number of sites on the genome. In early G1 (i.e., in small daughter cells), these sites are filled with Whi5-inhibited SBF complexes to which Cln3 can bind tightly in a stoichiometric fashion. Once bound, Cln3 slowly hypo-phosphorylates Whi5 and dissociates in the process.

**Figure 4.**
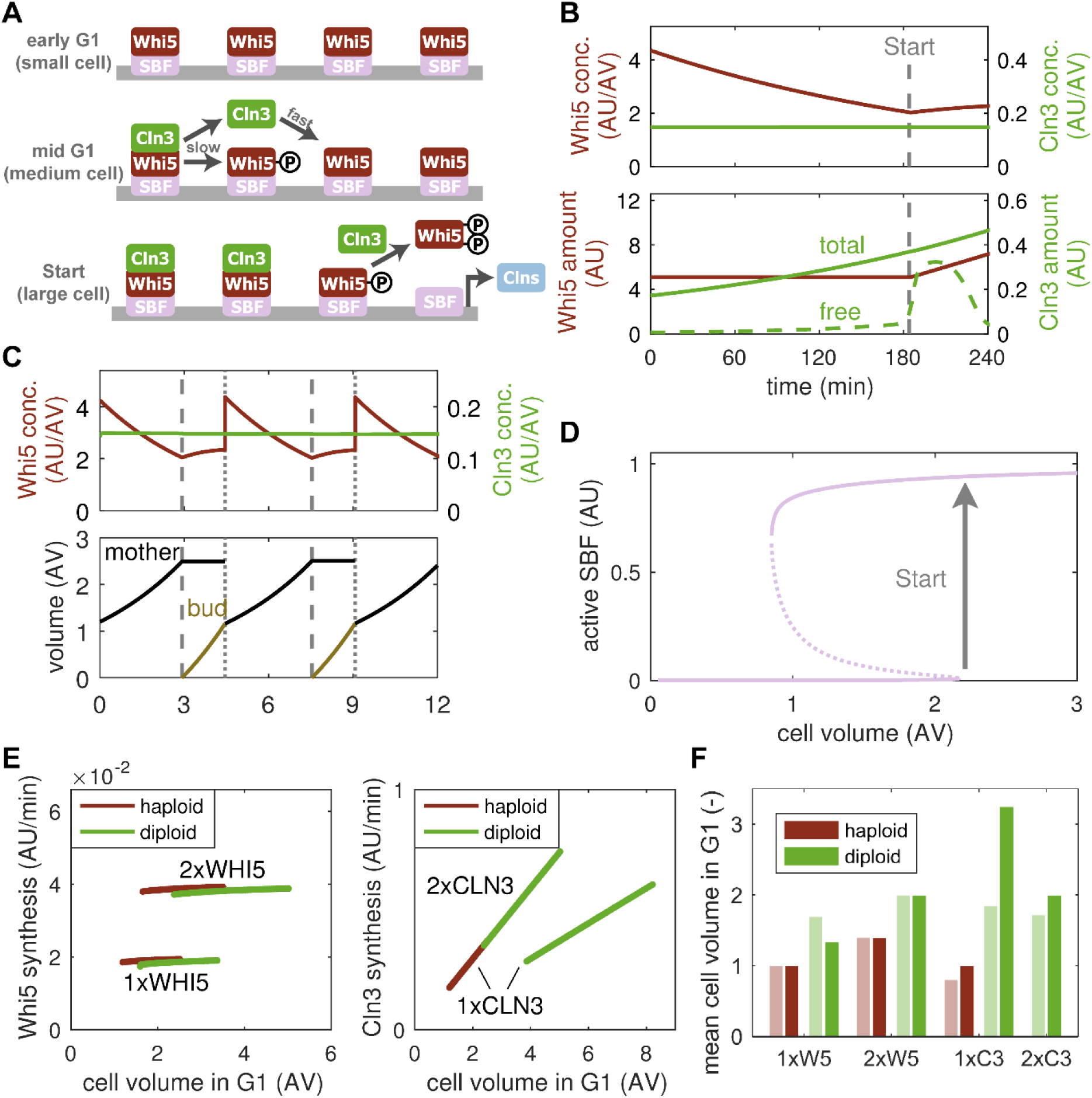
Titration-of-nudear-sites model captures ploidy effects on cell size. (**A**) Schematic of the titration model. SBF occupies a fixed number of sites on the genome and is inhibited by Whi5 in early G1. Gradually accumulating Cln3:Cdk1 binds to and slowly hypo-phosphorylates Whi5. When all sites are filled (or hypo-phosphorylated), free Cln3 emerges and hyper-phosphorylates Whi5, liberating SBF. (**B**) Total concentration (top) and amount (bottom) of Whi5 and Cln3 in a growing cell. The amount of free Cln3 is indicated in the bottom panel (dashed green line). Vertical dashed line marks START. (**C**) Concentration of cell cycle regulators (top) and cell volume (bottom) over multiple generations. Dashed and dotted vertical lines mark Start and division, respectively. Model follows the daughter cell (bud) after each division. (**D**) Stable (solid) and unstable (dashed) steady states of active SBF with respect to cell volume in the titration model. Arrow indicates START transition. (**E**) Simulation of Whi5 and Cln3 synthesis rates in haploid and diploid cells with the indicated copy number of *WHI5* and *CLN3.* (**F**) Mean cell volume in G1 for data in [1] (light bars) and simulations in D (dark bars). Values were normalized to haploid cells with one *WHI5* copy for each case.

However, it can rapidly rebind to unphosphorylated SBF:Whi5 (Fig. S1B). As the cell grows larger, the number of Cln3 molecules per cell increases (Cln3 is a size-dependent protein, whose concentration is maintained constant in G1) (Fig. 4B). This leads to a gradual accumulation of Cln3:Cdk1 heterodimers on Promoter:SBF:Whi5 complexes until all sites are filled, at which point free Cln3:Cdk1 kinase complexes emerge in the nucleus. Free Cln3:Cdk1 then promotes rapid hyper-phosphorylation of SBF-bound and free Whi5, facilitating the START transition (Fig. 4B). Similar to the inhibitor-dilution model, the titration model readily yields size homeostasis in consecutive generations (Fig. 4C) by coupling the passage through START to cell size (Fig. 4D).

When simulating changes in gene copy number, we observe that, similar to inhibitor dilution, the titration model correctly predicts protein synthesis rates (Fig. 4E). However, the titration mechanism also captures the increase in size between haploid and diploid cells with the same number of *WHI5* copies (Fig. 4E). In particular, diploid cells harbour twice the number of SBF binding sites, which require a higher amount of Cln3, and therefore a larger cell size, to be filled (Fig. S3A). Note that our model overestimates the size of diploid cells with one copy of *CLN3*(Fig. 4F). The cause for this discrepancy is that the absence of a second *CLN3* copy in diploid cells only reduces Cln3 synthesis rate by ∼15% (compare diploid cells with 1xCLN3 and 2xCLN3 in Fig. 3A, right panel), whereas the model predicts a reduction by ∼50% (Fig. 4E, right panel). After accounting for this, cell size predictions are much more accurate (Figs. S3B and S3C). It is not yet clear why a single *CLN3* can partially compensate for the second copy’s expression rate in diploid cells.

Further experimental evidence for a titration mechanism comes from an observed increase in cell size upon transformation of otherwise wild-type cells with a high copy number plasmid containing perfect SBF binding sites [20]. These decoy sites were proposed to change the size threshold for START by binding Cln3 such that an increased number of Cln3 molecules, and hence a larger cell size, is required to initiate the transition. Simulating this setup, our model does indeed show such an increase in size (Fig. S3D), providing further support for the existence a titration mechanism.

In summary, a combination of Whi5 dilution and Cln3 titration against SBF binding sites is not only able to capture protein synthesis rates but also the size of *WHI5*- and *CLN3*-mutant haploid and diploid cells and of cells harbouring an increased number of SBF binding sites.

### Sizer and timer combine to yield a phenomenological adder

Historically, three different strategies have been proposed to maintain cell size homeostasis: the sizer, where a cell cycle transition is triggered once the cells reaches a critical target size; the timer, whereby the cell cycle takes a constant amount of time; and the adder, postulating that cells add a constant volume each generation [25,26]. Each of these concepts may apply to the complete cell cycle or only to a certain cell cycle phase, and all of them generate characteristic size patterns that can be probed experimentally (Fig. 5A). An ideal sizer mechanism suggests that the final volume at the end of the sizer period is independent of the initial volume, such that the added volume shows a linear slope of minus one, i.e., small cells need to grow more to reach the critical size. By contrast, exponentially growing cells that employ a perfect timer show a slope of plus one in the added volume as small cells grow less during the same time increment. Note that a slope of exactly one is only observed if cells double their mass within the phase that uses a timer, e.g. if the timer is employed over the whole cell cycle of a symmetrically dividing cell. Finally, an adder leads to a slope of zero since the added volume is assumed to be constant. We wanted to understand how these concepts connect to the mechanistic model of cell cycle control presented above.

**Figure 5.**
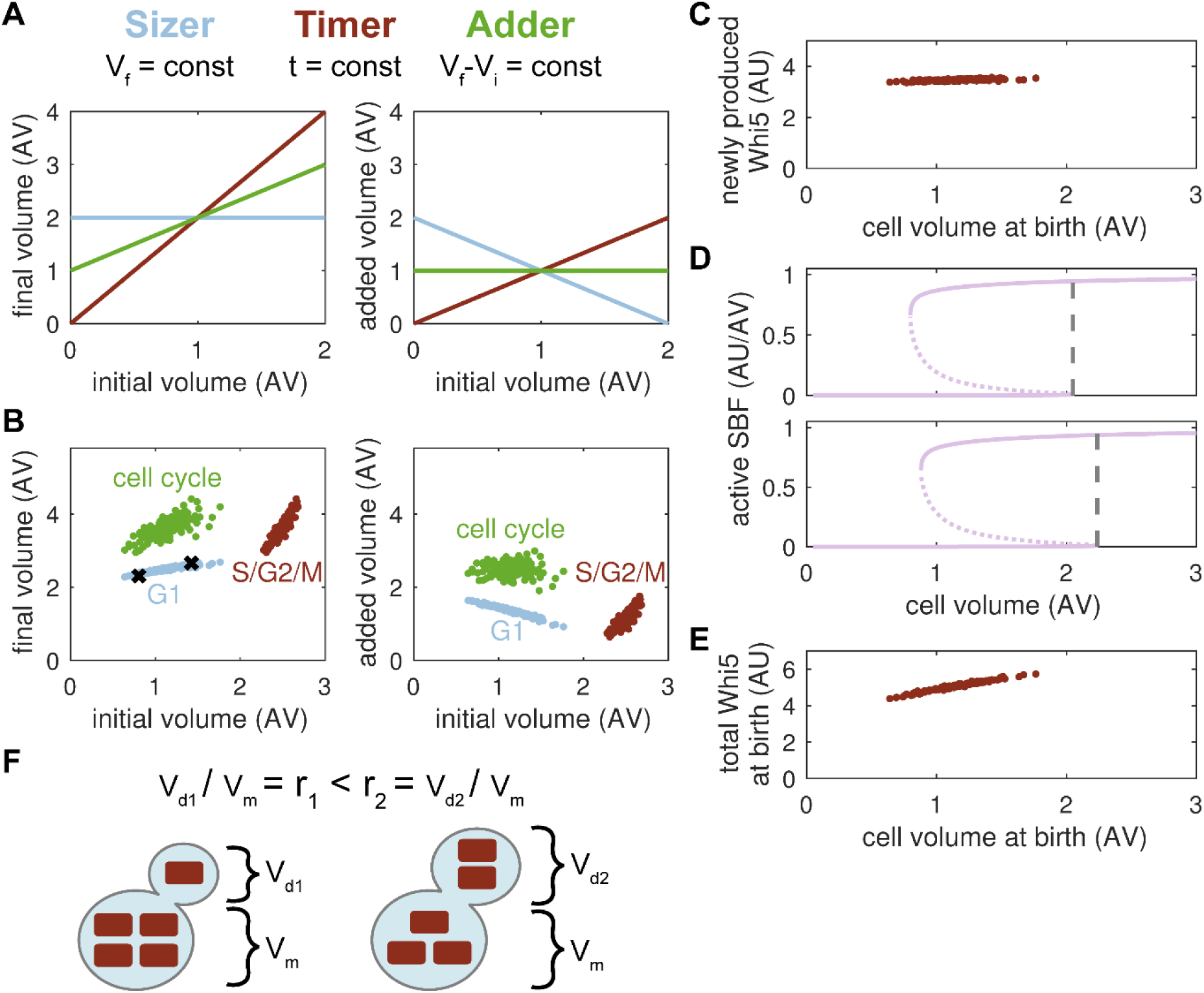
G1 sizer and S/G2/M timer combine to yield adder. (**A**) Theoretical predictions of the final volume (V_f_) and the added volume (V_f_ - V_i_) over the initial volume (V_i_) for an ideal sizer, timer and adder. Simulations assume exponential growth and, for the timer, a doubling of cell mass. (**B**) Simulation of the final volume and added volume over the initial volume for G1-phase, S/G2/M-phases and the whole cell cycle in the titration model. Black crosses mark cells shown in D. (**C**) Amount of newly synthesised Whi5 in cells of different birth size. (**D**) Stable (solid) and unstable (dashed) steady states of active SBF with respect to cell volume for the two cells marked with black crosses in B. Dashed lines indicate size threshold for START. (**E**) Total amount of Whi5 at cell birth for cells of different birth size. (**F**) Scheme of Whi5 distribution at cell division. Whi5 that is not bound to the genome, i.e., which diffuses freely through the nucleus or cytoplasm (red rectangles), is distributed according to the volume ratio of mother and daughter cell (r), which causes larger daughter cells to inherit more Whi5. V_d_, daughter cell volume; V_m_, mother cell volume.

Simulations of our titration model reveal that Gl-phase behaves like an imperfect sizer with smaller cells adding more volume during G1 (slope of −0.64; Fig. 5B, right panel) and cell size at S-phase entry showing a slight positive correlation with birth size (Fig. 5B, left panel). S/G2/M-phases, by contrast, exhibit a timer (see also Fig. S4A). The combination of a mechanistic sizer and a mechanistic timer yields a phenomenological adder with the added volume being virtually independent of cell size at birth (R = −0.02; Fig. 5B, right panel).

However, the added volume is not directly sensed by the system in any way. Instead the negative slope of the sizer compensates for the positive slope of the timer.

The results above raise the question as to why cells employ two seemingly different strategies in G1 and S/G2/M-phases, a sizer and a timer, respectively. Presumably, S, G2 and M-phase are completed fast, with a size-independent timing, to allow the mother cell to start the next budding event, while size control is relegated to the daughter cell’s G1 phase. In addition, a timer period of constant length in combination with a size-independent Whi5 synthesis allows the cell to produce a constant, size-independent amount of Whi5 per cell cycle (Figs. 5C and 2E). This constant Whi5 amount is part of the mechanism that tunes G1 length with respect to birth size. Hence, the S/G2/M timer helps to set up the G1 sizer. We also note that our simulations show an imperfect sizer with a slight positive correlation between the cell volume at START and the birth volume (Fig. 5B, left panel), as has been found experimentally [21,25]. Whereas an ideal sizer requires the size threshold for START to be independent of birth size, we find that cells which are larger at birth progress through START at a slightly larger size (Figs. 5B and 5D). According to our model, the main reason for this threshold change is the distribution of Whi5 molecules at cell division. In particular, larger cells are born with slightly higher amounts of Whi5 (Fig. 5E), since some of the Whi5-containing complexes are distributed according to the volume ratio of mother and daughter cell (Fig. 5F). It is primarily by this mechanism that birth size affects the size threshold for START in our model, as shown in Fig. S4B, where we manually set the Whi5 amount at birth to a constant value (birth size-independent) and find that the model behaves as an almost ideal G1 sizer.

In summary, our model shows that size control in budding yeast uses an S/G2/M timer that helps to produce a constant amount of Whi5 per cell cycle and to facilitate a sizer in the G1 phase of daughter cells. Both mechanism combine to yield a phenomenological adder over the whole cell cycle. However, the size-dependent distribution of Whi5 at cell division can cause an imperfect adjustment to size differences at birth.

## Discussion

Balanced growth, achieved by coupling cell division to the increase in cell mass, is crucial to cell survival as progressive changes in size over generations would eventually lead to a breakdown of biochemical processes. In this study, we developed a mechanistic mathematical model for size control in budding yeast based on the differential expression of cell cycle regulators in growing cells. We show that the interplay of size-dependent and size-independent synthesis of these regulators can establish a size threshold at START and facilitate size homeostasis.

It has long been recognised that the amount of most proteins in a cell increases with cell size [27,28], such that protein concentrations remain constant and reaction rates are unaffected by growth [12]. This has also been observed for the majority of cellular mRNAs, suggesting that adaptation to volume growth occurs at the transcriptional level [22,27,29–31]. Based on these observations, we propose a general mathematical model for gene expression in growing cells which assumes that a limiting component of the transcription machinery, which we named TM and that may correspond to an RNA polymerase or factors influencing chromatin accessibility [18], is produced in an autocatalytic manner by transcribing its own mRNA. Under conditions where nutrients and precursors are not limiting, this leads to an exponential increase in TM. If we assume that the increase in cell volume depends on proteins that are themselves transcribed by TM, the exponential rise in TM directly translates into an exponential increase in cell volume and it naturally leads to a direct proportionality between both, i.e., protein synthesis rates per unit volume are constant. This scaling is an emergent property of the system and does not require complex regulation or a dedicated mechanism that measures size and tunes transcriptional capacity accordingly. In very large cells, genes become saturated, at which point transcription rates remain constant and cell growth transitions into a phase of linear increase. These features of the model are consistent with a large body of experimental literature showing exponential growth of cell volume and transcription for small cells which plateaus when cells exceed a certain size [12,21,22,25,32].

Given this model of gene expression, two different types of genes emerge in our simulations based on their affinity to TM. Genes that bind TM with high affinity are saturated early, in small cells, and thus show size-independent protein synthesis. Consequently, they give rise to size-independent proteins, whose amount is constant, leading to a decreasing concentration in growing cells. Whi5 is an example of such a protein [13]. Due to their high affinity, size-independent genes compete efficiently for TM and an increase in their copy number, due to gene or genome duplication, directly translates into an increased synthesis and concentration. We hence propose that, in the context of size control, size-independent genes can act as gene-copy-number sensors. Beyond size regulation, the genes might encode proteins that need to be present in a fixed proportion to the genome content, e.g., transcription factors or histones. By contrast, size-dependent genes bind TM with lower affinity, such that their occupation by TM increases proportional to cell volume. Through this mechanism, their proteins can maintain a constant concentration until the gene is saturated. We propose that the majority of proteins uses this type of control, Cln3 being a concrete example [13]. Due to their characteristics, size-dependent genes share TM among themselves, such that an overall ploidy increase does not result in an increase in protein concentration. Size-dependent genes can hence act as sensors for the copy-number-to-ploidy ratio, the gene dosage. Variations of the affinity constants between the two extremes may lead to intermediate expression patterns, including genes that can switch from size-dependent to size-independent expression within a given range of cell sizes.

By incorporating the gene expression model into a model of the yeast cell cycle, we show that size-independent synthesis of the inhibitor Whi5 and size-dependent synthesis of the activator Cln3, a mechanism termed inhibitor dilution [13], can indeed establish size control at START. It is important to note that, because Whi5 is a stoichiometric inhibitor of SBF without catalytic activity [15,16], we have assumed in our inhibitor-dilution mechanism that Whi5 and SBF form a tightly bound complex. In addition, we assumed that phosphorylation of Whi5 by Cln3 breaks up the complex and liberates SBF. Considering that SBF maintains a constant concentration, as has been shown experimentally for one of its subunits [13], Whi5 is therefore in fact countered by two size-dependent activators, Cln3 and SBF. Given these molecular interactions, our results suggest that, in the inhibitor-dilution paradigm, the rising number of SBF molecules in a growing cell eventually overcomes inhibition by exceeding the constant number of Whi5 molecules (see Supplementary Text for details). Cln3 merely sets the threshold at which SBF activation occurs by keeping a fraction of Whi5 molecules in a phosphorylated (inactive) state. Because this fraction does not change appreciably with cell size, Cln3 is not directly involved in creating the size-dependent signal that facilitates START in our version of the inhibitor-dilution model. Hence, between the inhibitor-dilution model and the titration-of-nuclear-sites mechanism there exists an intriguing symmetry, in which Whi5 and nuclear sites are very much alike. Both are constant in number and proportional to DNA content and both titrate away an activator. We also show that the gradual increase in SBF activity in response to cell volume growth that is caused by Whi5 dilution is converted into an all-or-nothing decision by a bistable switch located at START. This switch is created by a positive feedback loop on SBF activity and it establishes a strict size threshold of START. Hence, positive feedback and bistability are used to implement a size checkpoint in G1.

While inhibitor dilution is able to maintain size homeostasis and reproduce the size increase seen in diploid cells, it fails to explain why an increase in ploidy at a constant number of *WHI5* copies leads to larger cells. Such a change does not alter the expression of Whi5 and Cln3 and should hence not affect the size at START. Even the delay in S/G2/M progression observed experimentally [13] is unable to reproduce these size changes in our model, suggesting that ploidy influences cell size beyond an effect through Whi5 and Cln3 expression and S/G2/M duration. Such an effect could be mediated by an as-yet-unknown inhibitor of START which is produced in a size-independent manner similar to Whi5. In this case, an increased expression of this inhibitor in diploid cells, due to a higher copy number of its gene, would cause the observed size increase. However, a much more appealing hypothesis is that the genome itself acts as an inhibitor of START. In particular, the binding of SBF to a limited number of genomic sites, which was proposed based on experiments [20], essentially converts SBF into a variable that does not change in number with cell size, as only the SBF that is bound to the genome would affect the START transition. Since the number of Whi5 molecules is constant as well, the amount of Whi5:SBF complexes, assuming tight binding between both, is not changing with cell size. However, the number of Cln3 molecules increases, such that Cln3 titrates against Whi5:SBF complexes on the genome. At a particular threshold size, Cln3 exceeds the number of Whi5:SBF complexes, leading to a sharp increase in free Cln3 that can trigger START. Positive feedback is again used to convert this increase into an all-or-none decision. In this context, a diploid cell is larger because it contains twice the number of SBF binding sites, requiring more Cln3 molecules to trigger START. Hence, the genome itself, through providing SBF binding sites that titrate Cln3, acts as a START inhibitor, using a form of distributed control (binding sites distributed throughout the genome) instead of a single gene product such as Whi5. We show that this Cln3 titration model is consistent with *WHI5* and *CLN3* mutants and experiments that express additional SBF binding sites, which lead to increased cell size at START [20]. Also note that a size-independent synthesis of Whi5 in the titration model is beneficial as an increasing Whi5 production in large cells would impair their progressing through START, compromising size control. Moreover, the proportional increase in Whi5 synthesis with gene copy number allows for a constant ratio between Whi5 and SBF molecules on binding sites in cells with increasing ploidy providing an intriguing hypothesis for why Whi5 is synthesised in a size-independent manner.

In recent years, studies of bacterial size control have argued for an adder-type mechanism, whereby cells add a constant increment of cell mass per cycle [33,34]. A similar type of behaviour was found between two budding events in *S. cerevisiae* [32]. Yet, it remained unclear whether cells actively sense the added mass and use this information to regulate cell cycle events, a scenario later referred to as a mechanistic adder [21]. From our simulations, we indeed observe the presence of an adder over the whole cell cycle, with no correlation between the added cell mass and the volume at birth. However, this behaviour does not result from a direct mechanism, but rather from a combination of a mechanistic sizer in G1 and a mechanistic timer in S/G2/M, which is in excellent agreement with a recent study arguing that the adder phenomenon emerges from independent pre- and post-START controls [21]. Similar to these and other experiments [5,21], our titration model shows an inverse proportionality between G1 length and birth size, and an imperfect sizer mechanism. We propose that adaptation is imperfect because of a volume-dependent distribution of Whi5. An ideal sizer, where START size is independent of birth size, requires that each daughter cell receives a constant amount of Whi5. However, Whi5 complexes that diffuse freely in the nucleus or cytoplasm would be distributed based on the size of the daughter cell, with large cells receiving a larger increment of Whi5 that keeps them in G1 for longer. In our model, this results in a weak birth-size dependence of the START threshold and imperfect size control. This might be one reasons why cells do not rely on a pure inhibitor-dilution mechanism, which would exacerbate the influence of Whi5 distribution, but instead use a combination of Whi5 dilution and Cln3 titration. In addition, Cln3 is a highly unstable protein [35], and thus provides a snapshot of the current transcriptional capacity and cell volume, while Whi5 was produced in the previous cycle, inevitably including some form of memory of past growth conditions.

In summary, our study provides a mechanistic model of gene expression and cell cycle regulation in budding yeast that readily shows size homeostasis. Since the control network of START in budding yeast is structurally similar to restriction point control in mammalian cells, similar mechanisms could be at work during mammalian size control.

## Acknowledgements

FSH and BN are funded by the BBSRC Strategic LoLa grant (BB/M00354X/1). JJT acknowledges financial support from the US National Institutes of Health (5R01-GM078989-12) administered through Colorado State University (PI: Jean Peccoud).

## Author contributions

FSH, JJT, BN: designed the research, developed the models, analysed the data and wrote the paper. RL: developed and analysed an early version of the inhibitor-dilution model. JJT, BN: secured funding.

## Materials and Methods

### Mathematical modelling

Our models for budding yeast size control comprise sets of ordinary differential equations (ODEs). These ODEs describe the dynamics of genes and proteins in terms of their molecule number rather than concentration, which is used by most biochemical models that do not account for cell volume growth. In the following, we explain each of the two models (inhibitor dilution and titration of nuclear sites) in detail, starting with a generic description of gene expression that underlies both models.

#### Gene expression

Gene expression was modelled with the aim to capture the size-dependent and size-independent synthesis of proteins, which causes them to maintain their concentration or become diluted by cell growth, respectively. As these two types of regulation occurs at the transcriptional level [12,22], we accounted for the amount of transcription machinery (*TM*). In our model, TM can bind to size-independent (*GI*) and size-dependent (*GD*) genes, leading to the formation of TM-gene complexes (*GITM* and *GDTM*, respectively).

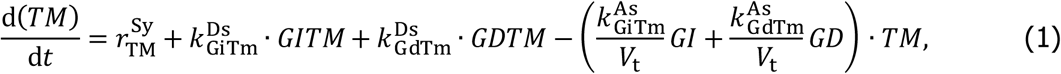

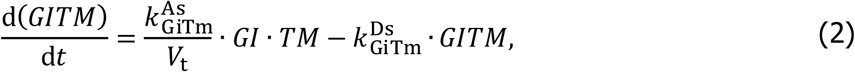

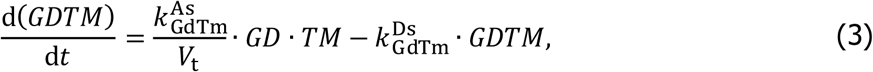

where association rates are denoted 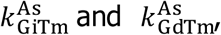 and dissociation rates are 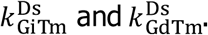 The main difference between both gene types is their binding affinity, i.e., size-independent genes bind more tightly to TM 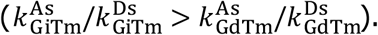 Furthermore, we assumed that TM is stable and synthesised with rate 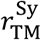 (see Eq. 5). Note that the rate of a bimolecular (binding) reaction is inversely proportional to the total volume (*V*_t_) of the system, reflecting the fact that two molecules have a harder time finding each other inside a larger volume [36]. This volume may increase according to

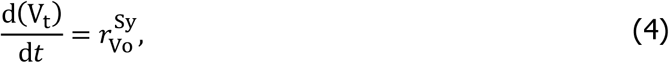

with the volume synthesis rate 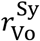 (see Eq. 6). Since the amount of most proteins in a cell increases with cell size [12,27,28], i.e., their synthesis is size-dependent, and proteins themselves are directly or indirectly responsible for cell volume growth, we assumed that both the synthesis rate of TM and cell volume are proportional to the number of transcriptionally active size-dependent genes (*GDTM*).

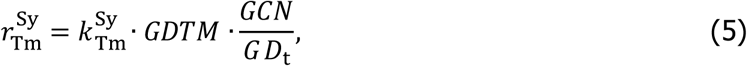

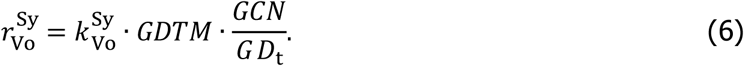

Here, 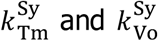 are the rate constants for TM and volume synthesis, respectively. Note that in a haploid cell only one of all size-dependent genes (*GD*_t_) would be expected to encode for a specific protein, which is why we scale the synthesis rates using the gene copy number (*GCN*). Following this scheme, size-independent (*P*_i_) and size-dependent (*P*_d_) proteins accumulate according to

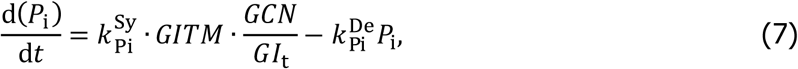

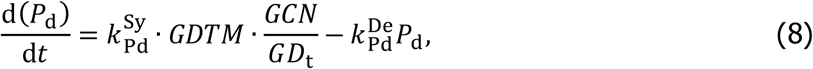

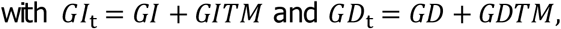

where 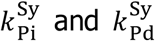 are rate constants for protein synthesis, and protein degradation rates are denoted 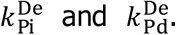 Conservation equations hold for the total number of size-independent and size-dependent genes (*GI*_t_ and *GD*_t_, respectivey). This generic model for gene expression is used in the following to describe budding yeast size control in both the inhibitor-dilution and nuclear-sites-titration models.

#### Inhibitor-dilution model

Using the principles of gene expression shown above (Eqs. 1-6), we developed a model of the yeast cell cycle that accounts for cell volume growth and size control through inhibitor dilution. In particular, the volume of the mother cell (*V*_m_) and the daughter cell (*V*_d_), i.e., the bud, grow according to

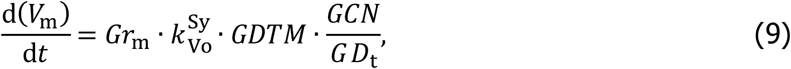

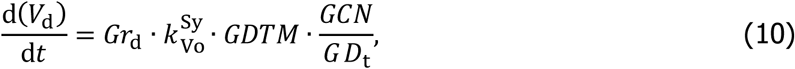

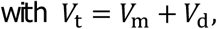

where *Gr*_m_ and *Gr*_d_ are binary variables that control whether growth is directed into the mother or daughter. These variables were introduced based on experimental observations suggesting that, during the budded period, volume growth primarily occurs in the bud and not the parent cell [4]. Note that during the budded period, the total cell volume (*V*_t_) comprises the volume of both mother and daughter.

In our model, cell cycle progression occurs through the expression of three types of cyclins: the G1 cyclins Cln3 (*CLN3*) and Cln1/2 (*CLN*), and the mitotic cyclins Clb1/2 (*CLB*).

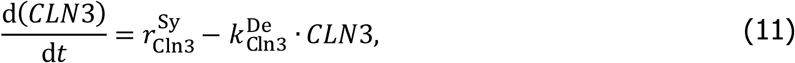

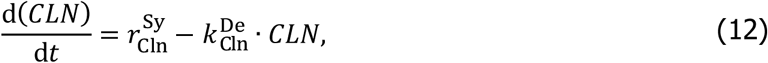

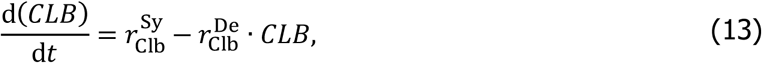

with 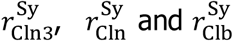 denoting synthesis rates, and 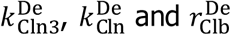 degradation rates. These rates were defined as follows:

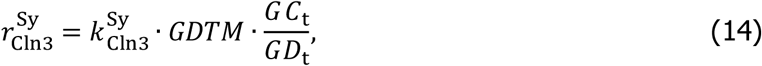

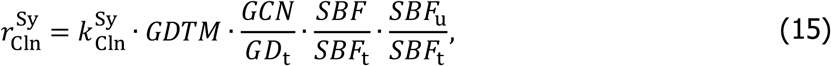

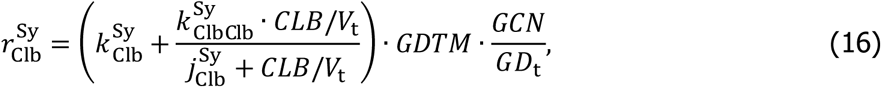

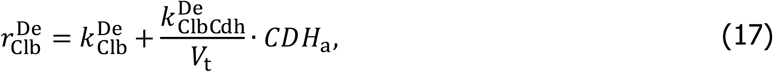

where 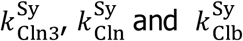 are the rate constants for constitutive synthesis, and 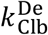 the rate constant for constitutive Clb1/2 degradation. All three proteins were assumed to be size-dependent, as are the majority of cellular proteins [12,27,28]. Note that we introduced a parameter for the copy number of the *CLN3* gene (*GC*_t_) to vary it independently from overall cell ploidy (*GCN*). Since expression of Cln 1/2 depends on the SBF transcription factor [37], we scaled the Cln synthesis rate by the fraction of free (not inhibited by Whi5; see Eq. 20) SBF (*SBF*) to total SBF (*SBF*_t_). Furthermore, SBF can become inhibited through phosphorylation by B-type cyclins in the later stages of the cell cycle [38]. Hence, Cln1/2 synthesis is also scaled by the fraction of unphosphorylated SBF (*SBF*_u_; Eq. 22). For Clb1/2, we assumed that, in addition to constitutive production 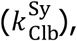 there is auto-activation [38] with maximal rate 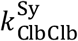 and a Michaelis-type constant 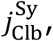 which causes saturation of the rate at high Clb1/2 levels. Note that auto-activation in this equation depends on the concentration of Clb1/2; hence, we divided by the total cell volume (*V*_t_). Finally, the degradation of Clb1/2 depends on a constitutive rate 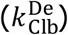 and on degradation by active APC/C^Cdh1^ (*CDH*_a_) with rate constant 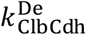 in a concentration-dependent manner [39].

In the inhibitor-dilution model, Whi5 (*WHI*) and free SBF (*SBF*) bind in a concentration-based manner to form Whi5:SBF complexes (*WHISBF*) that are devoid of activity, while phosphorylated Whi5 (*WHI*_p_) is unable to inhibit SBF [15,16].

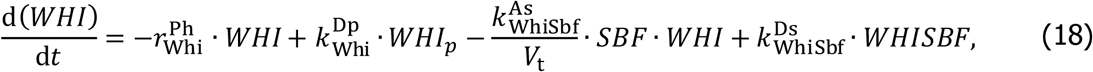

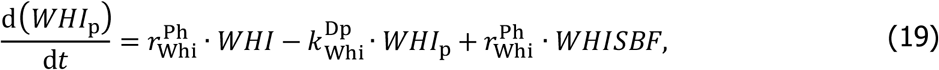

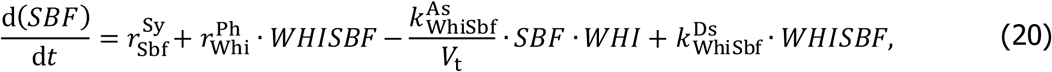

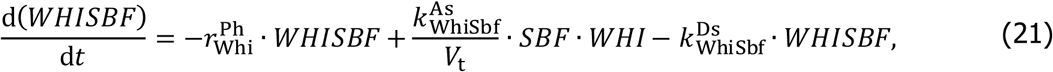

where assocaton and dissociation rates for Whi5 and SBF are denoted 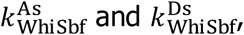 respectively. The phosphorylaton and dephosphorylation rates of Whi5 are 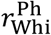 (see Eq. 26) and 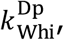 respectively. Note that the phosphorylation of Whi5 in Whi5:SBF complexes leads to their dissociation, which activates SBF. SBF is synthesised with rate 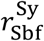 (see Eq. 25).

We assumed that the inhibitory phosphorylation of SBF is independent of its binding status and hence treated the state variables of phosphorylated (*SBF*_p_) and unphosphorylated (*SBF*_u_) SBF independent from the SBF variables shown above.

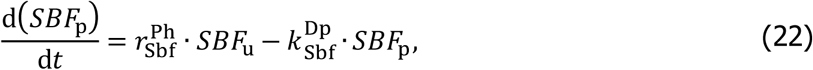

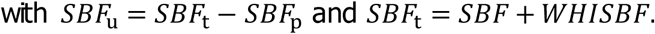

Here, 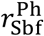 (see Eq. 27) and 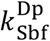 represent inhibitory phosphorylation and dephosphorylation of SBF, respectively. Conservation equations hold for the total amount of SBF (*SBF*_t_).

Production of Whi5 is restricted to S/G2/M phases, i.e., the budded period, and daughter cells receive a larger proportion of Whi5 at cell division [13]. Hence, we introduced newly produced Whi5 (*WHI*_n_) and tied its synthesis to the binary variable for bud growth (*Gr*_d_).

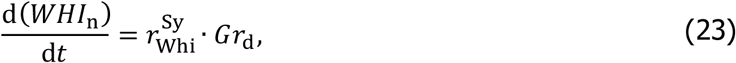

where the rate of Whi5 synthesis is denoted 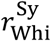 (see Eq. 24).

The synthesis and phosphorylation rates for the equations above were defined as follows:

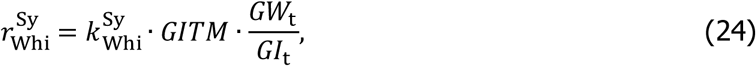

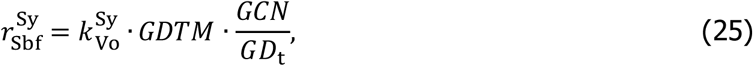

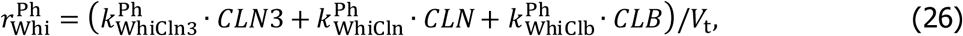

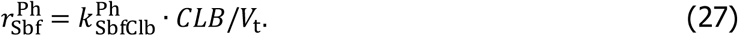

Here, Whi5 synthesis occurs with rate constant 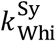 from a size-independent gene [13] with copy number *GW*_t_, which we introduced to vary the number of *WHI5* copies independently of ploidy. Whi5 is the only size-independent protein in the model (*GI*_t_ = *GW*_t_). Without loss of generality, SBF synthesis was assumed to occur with the same rate as cell volume synthesis 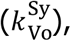 such that the SBF concentration remains constant at 1 AU/AV. In our model, Whi5 is phosphorylated by Cln3, Cln 1/2 and Clb1/2 [15–17] with rates 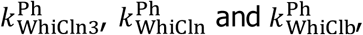 respectively, in a concentration-based manner. Similarly, SBF is phosphorylated by Clb1/2 with rate 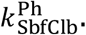

In order to exit the cell cycle, cells need to degrade cyclins using the APC/C. Our model accounts for two of its forms: APC/C^C dh1^ (*CDH*) and APC/C^C dc20^ (*CDC*), both of which can be present in an active (subscript a) and an inactive (subscript i) configuration.

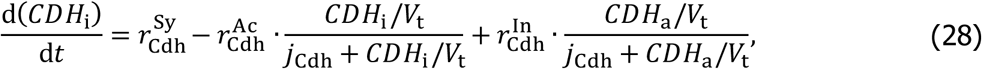

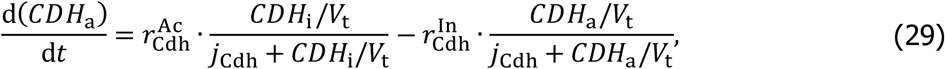

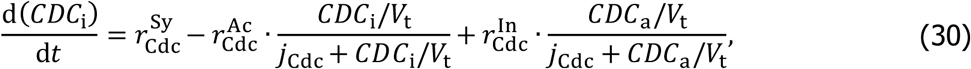

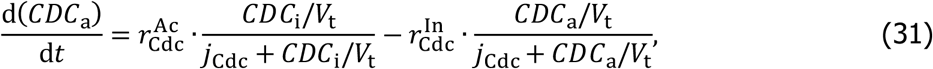

where the Cdh1 and Cdc20 subunits are synthesised with rates 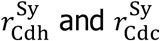 (see Eqs. 32-33), respectively, and binding of these subunits to the APC/C is assumed to be instantaneous and limited by subunit availability. Based on previous models [40], activation and inactivation of the APC/C is assumed to occur via concentration-dependent, Michaelis-Menten-type kinetics with rates *r*^Ac^ and *r*^In^, respectively. The corresponding Michaelis-Menten constants are denoted *j*_cdh_ and *j*_cdc_. The rates of subunit synthesis and APC/C activation and inactivation were defined as follows:

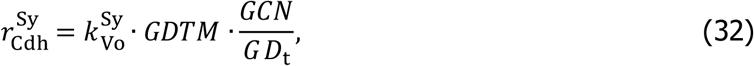

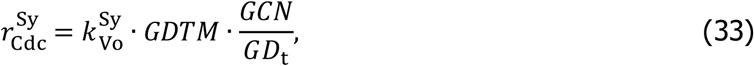

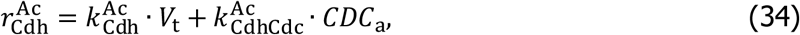

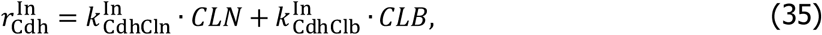

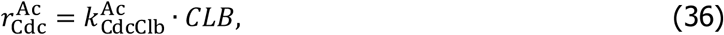

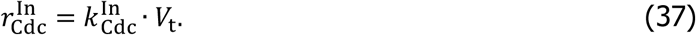

Here, synthesis of Cdh1 and Cdc20 occurs similar to SBF (described above) to a constant concentration of 1 AU/AV. APC/C^C dh1^ is activated with the constitutive rate 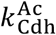 and by active APC/C^Cdc20^ with rate 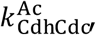 while inactivation is mediated by Cln 1/2 and Clb1/2 with rates 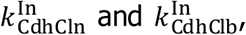 respectively. Similarly, APC/C^Cdc20^ is activated by Clb1/2 with rate 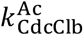 and inactivated with constitutive rate 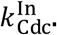

To account for genome replication and cell division, we introduced two events in our model. The first is triggered when Cln 1/2 increase above a threshold concentration, which induces bud growth and genome duplication.

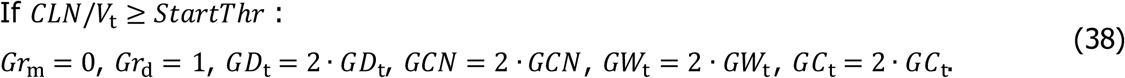

Similarly, cell division occurs when the combined concentration of Cln1/2 and Clb 1/2 falls below the threshold that maintains mitosis.

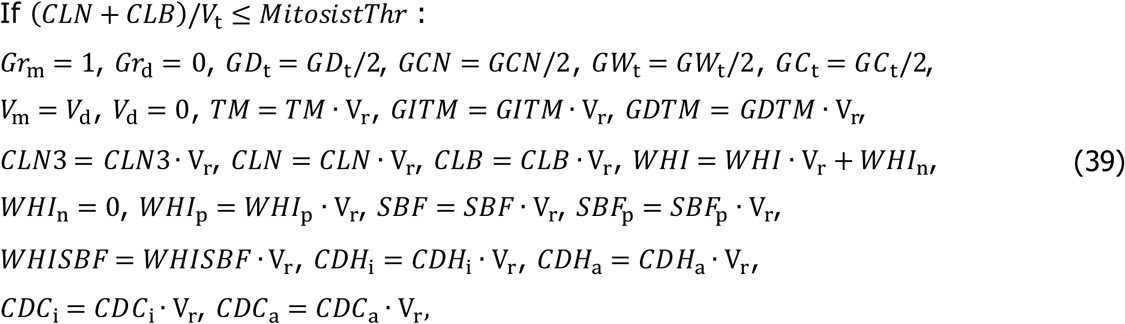

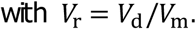

Note that we followed the daughter cell (bud) after budding as size control mainly takes place in these small, new-born cells [3–5]. Hence, at cell division, we assigned V_d_ to K_m_ (the daughter becomes the new ‘mother’ cell) and directed growth into the new cell. Moreover, the genome is split evenly, such that the new cell receives half of all gene-related variables. The remaining proteins are divided based on the volume ratio of mother and daughter cell before division (*V*_r_), assuming that these proteins can freely diffuse in the cytoplasm or nucleus. However, we assumed that all newly produced Whi5 is directed towards the daughter cell. This is consistent with experimental evidence showing a higher concentration of Whi5 in daughter cells compared to their mothers after division [13] and with the fact that mother cells exhibit a short G1 phase [4,5]. In particular, as mother cells do not increase significantly in volume after START, any newly produced Whi5 that remains in the mother cell would increase its Whi5 concentration above the previously passed threshold for START, thus extending the ‘old’ mother’s next G1-phase and leading to further mother-cell growth (our unpublished simulations). Also note that we assumed transcriptionally active genes rapidly adapt to the new concentration of TM and hence multiplied these variables by *V*_r_ as well.

#### Titration of nuclear sites model

Our titration model is based on the principles of gene expression outlined above and largely employs the same equations than the inhibitor-dilution model. The main differences between both models relate to the interaction of Cln3, Whi5 and SBF (see also *‘Differences between inhibitor-dilution and titration model* in the Supplement).

The growth in mother and daughter cell volume was modelled as described in Eqs. 9-10, with both depending on transcriptionally active size-dependent genes. Similarly, Cln 1/2 and Clb1/2 are synthesised from such genes following Eqs. 12-13 and 15-17. We modelled the interaction of Cln3, Whi5 and SBF according to the titration hypothesis put forward by Wang et al. [20]. In particular, we assumed that there is limited number of nuclear sites (*NS*_t_) for SBF binding. For the sake of simplicity, we only accounted for the SBF that is bound to these sites and neglected freely diffusing SBF, such that all SBF-related variables refer to SBF on nuclear sites and total SBF (SBF_t_) is constant at

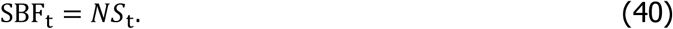

Cln3, Whi5 and SBF were assumed to interact in a two-step process (see Fig. S1B). First, Whi5 binds to SBF and forms Whi5:SBF complexes (*WHISBF*), which inhibits SBF activity. Subsequently, Cln3 can bind to form a trimeric Cln3:Whi5:SBF complex (*CLN3WHISBF*) in which Whi5 becomes hypo-phosphorylated, causing the dissociation of Cln3 and leaving a hypo-phosphorylated, but still inhibited, Whi5-P:SBF complex (*WHFpSBF*). This complex can either be dephosphorylated or hyper-phosphorylated by free Cln3, which liberates SBF and induces Cln 1/2 expression. Forming a postive feedback, Cln 1/2 promotes further hyper-phosphorylation of *WHIpSBF* and of free Whi5, preventing the re-inhibition of SBF. Following these considerations, the number of free Whi5 (*WHI*) and phosphorylated Whi5 (*WHI*_p_) molecules is given by

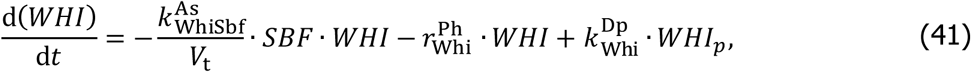

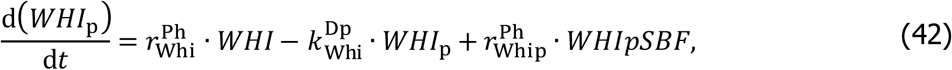

where 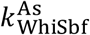 denotes the rate constant for Whi5-SBF binding and 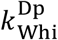 represents the dephosphorylation of Whi5. Free Whi5 is phosphorylated with rate 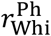 (see Eq. 47), white hypo-phosphorylated Whi5 in Whi5:SBF complexes is hyper-phosphorylated with rate 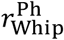 (see Eq. 48). Interaction of Cln3, Whi5 and SBF occurs according to

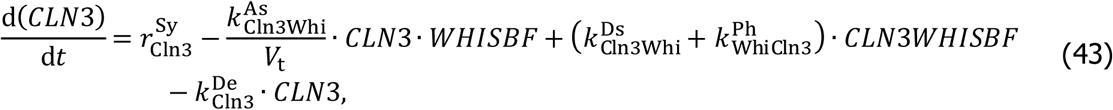

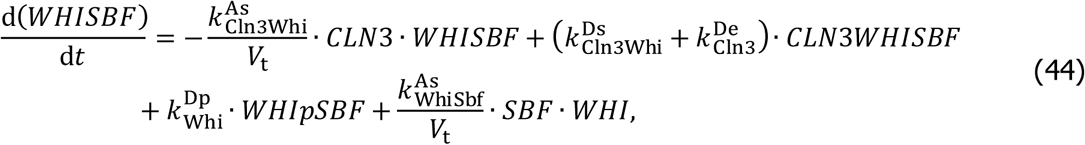

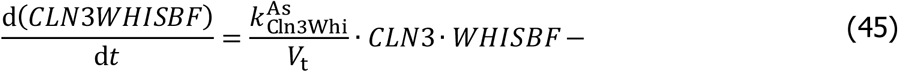

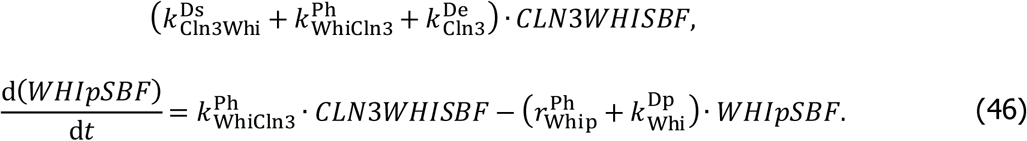

Here, Cln3 is synthesised and degraded with rate 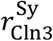 (see Eq. 14) and 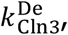 respectively, and it binds to and dissociates from Whi5:SBF complexes with rate 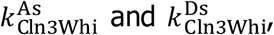 respectively. Hypo-phosphorylaton of Whi5 by Cln3 in the trimeric complexes occurs with rate constant 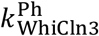 and leads to Cln3 dissociation. This phosphorylation can be reversed with rate or 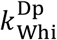 converted to hyper-phosphorylation with rate 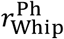 (see Eq. 48), which liberates SBF. The phosphorylation rates of Whi5 were defined as follows

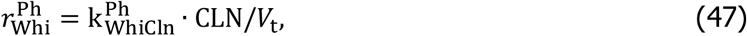

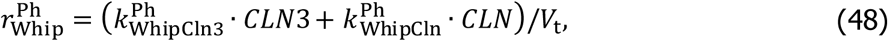

where the rate of free Whi5 phosphorylation by Cln1/2 is 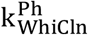 and the rates of hyperphosphorylation of *WHIpSBF* by Cln3 and Cln 1/2 are 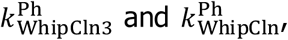 respectively. SBF on nuclear sites that is not inhibited by Whi5 (*SBF*) can be calculated from the conservation equation

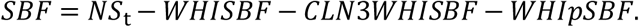

As in the inhibitor-dilution model, SBF activity is additionally regulated by an inhibitory phosphorylation (Eqs. 22 and 27), which is assumed to be independent of the other reaction steps SBF undergoes. Furthermore, the producton of Whi5 and the dynamics of APC/C^Cdh1^ and APC/C^Cdc20^ were modelled as described before (Eqs. 23-24 and 28-37).

Similarly to Eqs. 38-39, we introduced two events that represent START and cell division, respectively. An increase of the Cln 1/2 concentration above a threshold initiates bud growth and genome replication, which, in the titration model, includes an increase in the number of nuclear sites and SBF complexes bound to them:

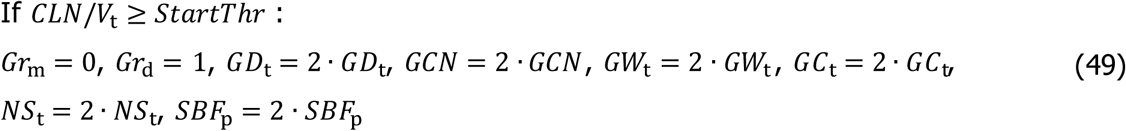

At cell division, which is initiated when the combined concentration of Cln1/2 and Clb 1/2 falls below the threshold that maintains mitosis, gene-related variables are divided equally between the two cells, while freely diffusing molecules are inherited based on the volume ratio of mother and daughter cell (*V*_r_).

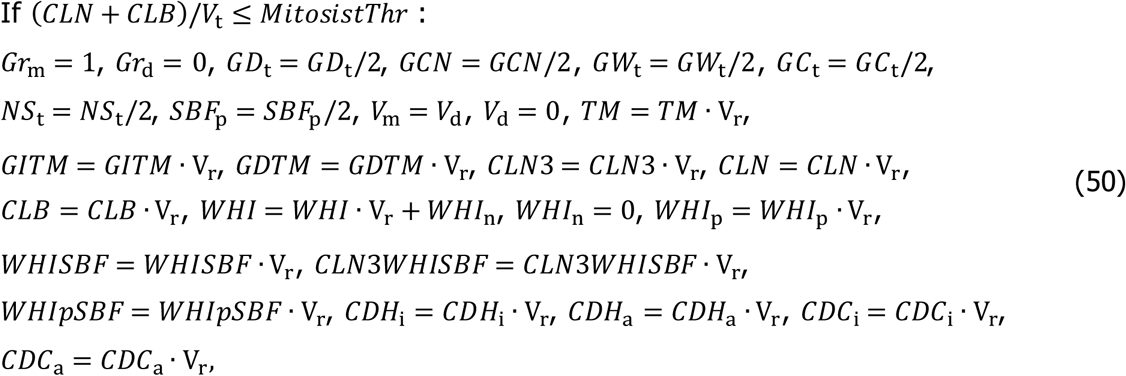

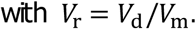

Again, the transcription machinery on genes is assumed to adjust rapidly to the new TM concentration, and so are the complexes of Whi5 and Cln3 with SBF on the nuclear sites.

#### Computation

Both size-control models were prepared in the Systems Biology Toolbox 2 [41] for MatLab (version 9.1.0 R2016b) and simulated with the CVODE routine [42]. Bifurcation diagrams were calculated using the freely available software XPP-Aut [43]. Models are provided as Files S1-3 in the Supplement and different versions are available at www.cellcycle.org.uk/publication. Model files were also deposited in BioModels [44] and assigned the identifiers MODEL1803220001 and MODEL1803220002. Parameter values and initial conditions are listed in Tables S1-4 and Table S5 shows the changes required to simulate ploidy mutants in Figs. 3, 4, S2 and S3.

In order to simulate cells of different sizes (e.g. in Fig. 2F and G), we varied the specific growth rate, with higher growth rates producing larger cells. In particular, the specific growth rate (*μ*) in our model follows from Eqs. 9-10 as

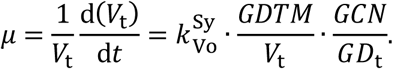

Since *GD*_t_ ≫ *GI*_t_ and almost all of the TM is bound to genes for the cell sizes we study here, the amount of transcriptionally active size-dependent genes can be approximated by the total number of TM (*GDTM* ≈ *TM*_t_). Moreover, we can calculate the transcriptional capacity per unit cell volume as

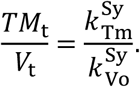

Taken together this gives

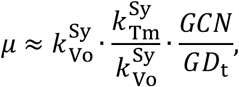

demonstrating that by changing both 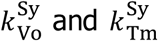 by the same factor, we can change the specific growth rate, while still maintaining the same transcriptional capacity per unit cell volume and thus similar protein expression. Accordingly, for simulations in Fig. 2F and G, 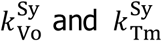 were multiplied by a factor *f* ∈ [0.75, 1.25]. For simulations in Fig. 5 and S4, we followed a single cell lineage over a large number of divisions to correlate cell sizes at different cell cycle stages. To obtain different cell sizes, we again varied the growth rate as described above, assuming that it changes at cell division. In particular, we assumed that the specific growth rate in the next cycle (*μ*_*n*+1_) is partly inherited from the mother cell’s growth rate (*μ_n_*) and partly influenced by stochasticity, e.g., by the random distribution of molecules at cell division, using

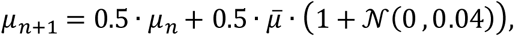

where *μ̄* is the average growth rate and 𝓝(0, *σ*) a normally distributed random variable with mean 0 and variance *σ*.

